# Filtering microbial populations with a magnetic field

**DOI:** 10.1101/2022.10.14.512298

**Authors:** Michelle Nazareth, Ece Kilinc, David Deamer

## Abstract

Magnetic fields strongly affect currents of electrically charged particles such as electrons, protons and other ions in solution. Because ionic currents of protons or sodium drive the rotation of bacterial flagella, it is possible that the motion of motile bacterial species will be affected if they swim through a strong magnetic field. We tested this prediction in mixed cultures of soil bacteria and observed that a magnetic field does in fact exert a filtering effect that alters the composition of the mixed population of motile species. We then monitored motility and growth to see if magnetic fields affected individual bacterial species (*Vibrio, Enterobacter sp*. and *Pseudomonas sp*.) The same magnetic field had no observable effect on motility or growth. Although magnetic fields may have served as a selective factor in the evolution of certain species such as magnetotactic motile bacteria, they do not appear to have a direct effect on the ionic current driving flagellar rotation.

## 1. Introduction

The majority of bacterial species are motile and use one or more flagella to move through an aqueous medium. To produce motility, each flagellum is driven by a flux of protons, or in some cases sodium ions, that pass through a motor embedded in the cell membrane [1]. The flux drives rotary motion of flagella and the rotation propels the bacterial cell at velocities ranging from 20 to 400 micrometers per second [2]. The flagella of *E. coli* rotate ∼100 times per second, and those of *Vibrio alginolyticus* approximately ten times faster [3]. *E. coli* flagella are driven by a flux of ∼1000 protons per rotation, or ∼100,000 protons per second [4].

A physical principle is that the motion of electrically charged particles is strongly affected if they are moving in a magnetic field. This is best known for currents of electrons either in a wire or a vacuum but is also true for any current of electrically charged particles. In a living motile bacterium there are three primary ionic species in motion: the proton or sodium ion current driving flagellar rotation, the protonic current through the ATP synthase and the electrical charges on flagellin, the protein that composes flagella.

Given a significant flux of approximately 10^5^ ionic charges per second through a flagellar motor, equivalent to a current of ∼20 femtoamperes, a strong magnetic field may have a measurable effect on bacterial motility. Furthermore, because different species of bacteria have a range of flagellar rotation rates there may be differential effects of a magnetic field on mixed populations of bacteria. This prediction was tested with populations of motile soil bacteria which were made to swim through a strong magnetic field of 1.25 teslas. We observed distinct differences of species composition compared to controls and conclude that a magnetic field has a filtering effect on certain species. However, when several individual species were exposed to a magnetic field no effect on motility or growth was observed. We conclude that the filtering effect of magnetic fields on mixed cultures is not due to a general effect on proton or sodium flux through the flagellar motor.

## 2. Materials and Methods

The primary goal was to determine whether there are any observable effects of a magnetic field on motile bacteria, so we began with mixed cultures of common soil bacteria. To this end, ∼ twenty grams of moist soil samples were collected and shaken with 200 mL of water. The soil-water mixture was passed through coarse paper filters to remove debris and large soil particles. Aliquots of the filtrate (0.5 mL) were added to 10 mL of 2% LB nutrient solution, then incubated for 18 hours at room temperature. The cultures were prepared at approximately weekly intervals throughout 2020 and 2021.

### Exposing motile bacteria to a magnetic field

Two holders were fabricated in which a 10 mL glass test tube could be held in place between the north and south poles of two neodymium magnets that produced ∼1.0 T magnetic fields across the diameter of the tube (Figure 1). In order to separate motile bacteria from sedentary species, 3.4 grams of sucrose were dissolved in 5 mL of the 18-hour soil bacterial culture to bring the sucrose concentration up to 2.0 M. The bacterial cultures with sucrose added were denser than LB medium so that 3 mL of fresh medium could be floated on top of 1.0 mL of the culture with added sucrose. Over an 18 hour period, motile species swam upward through the magnetic field into the fresh medium and accumulated on top while non-motile bacteria were left behind in the dense culture medium. Each experiment consisted of two tubes with magnets in place and two control tubes with no magnets present.

**Figure 1.**
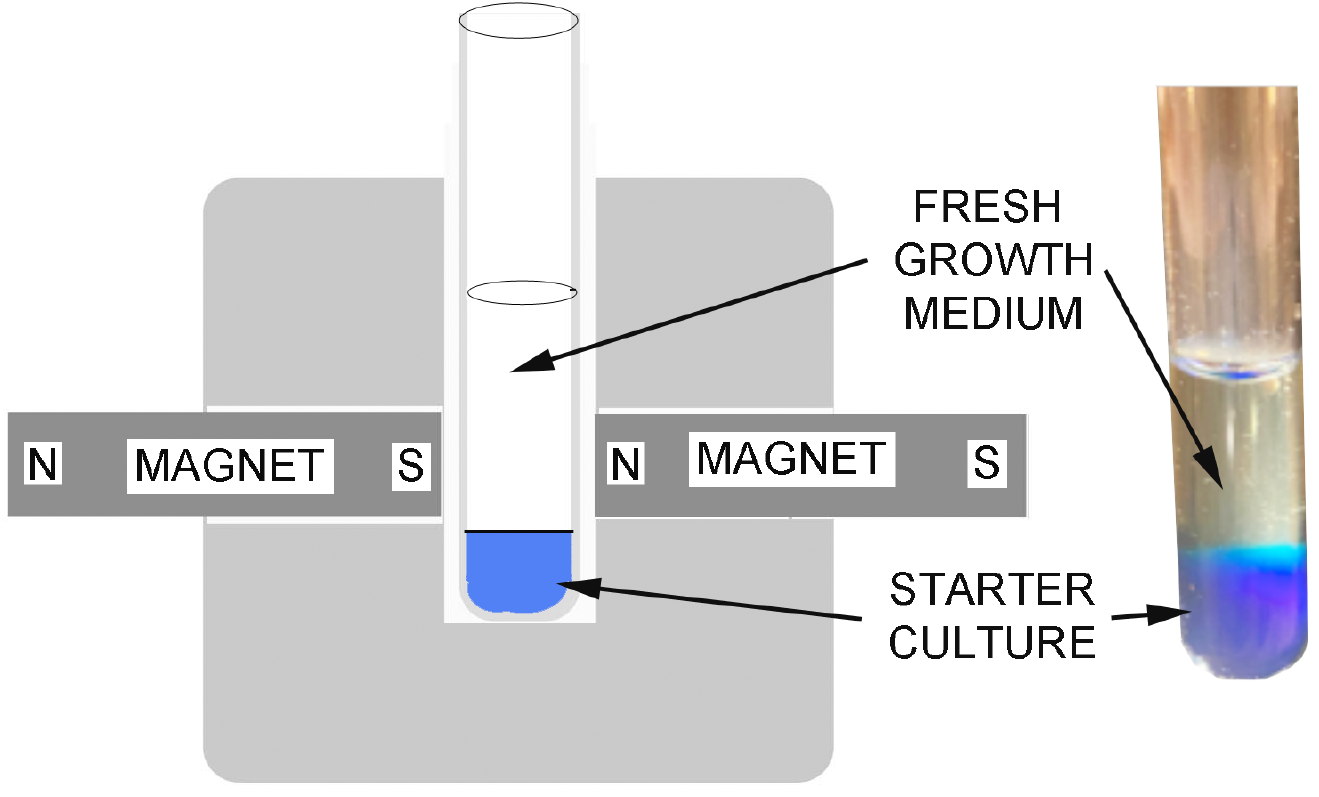
Experimental set up. Tubes were placed in wooden containers for 18 hours with or without the presence of a magnetic field. The photograph shows 1.0 mL of a bacterial LB culture mixed with enough sucrose to produce a dense 2.0 M solution. The upper phase is 3 mL of fresh LB medium that floats on top. Blue dye was added to indicate the demarcation between the starter culture and growth medium.

After 18 hours, the upper 0.5 mL of the solution containing motile bacteria was removed and 50 uL aliquots of the samples were Gram stained with a commercial kit and then viewed with a Zeiss Axiovert microscope. All photographs were taken at 1000X magnification using an oil immersion objective lens.

### 16S rRNA analysis

Samples for genomic analysis were run separately from those used for microscopy. After an overnight incubation, 0.5 mL aliquots from the top of control and magnetic samples were centrifuged to form a pellet. The pellets were preserved in ethanol and submitted to Creative BioGene: Microbiosci Inc. (Shirley NY 11967) where 16S rRNA analysis was performed to determine the species identities and relative numbers of species present in the control and magnetic samples. Figure 2 illustrates the preparation steps and bioinformatic analysis which were performed in two separate experiments in 2020 and 2021.

**Figure 2.**
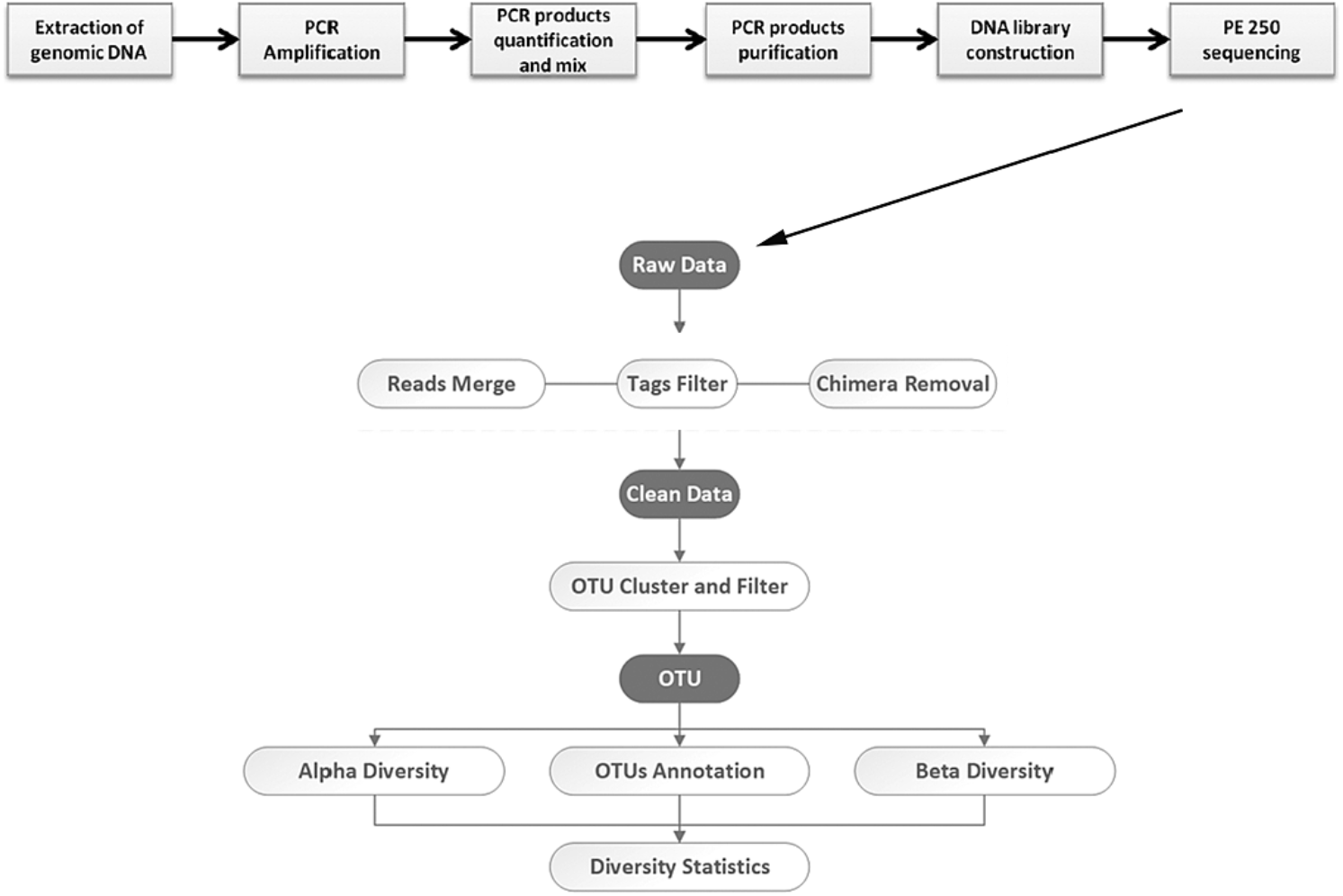
Method used by Creative BioGene to analyze bacterial pellets.

### Exposing known species of bacteria to magnetic fields

To test whether magnetic field affects motility and growth of known species of motile bacteria, two neodymium magnets 1 X 3 X 0.5 cm were glued lengthwise in slots milled into a square 20cm X 20cm wooden support. Aliquots of *Vibrio, Enterobacter* and *Pseudomonas* cultures were streaked heavily on a nutrient agar plate and incubated at 37° C overnight. The 0.3% agar motility medium contained 1.0 g yeast extract, 2.0 g NaCl, 2.0 g tryptone, and 0.6 g agar in 200 mL H_2_O. The pH was adjusted to 7.5 with dilute NaOH, then sterilized by autoclaving. The medium was used to prepare at least six petri plates for each bacterial species and allowed to gel overnight at room temperature.

To begin the motility experiments, single bacterial samples were isolated from the streak plate and stabbed in the center of each motility plate. The test plates were placed so that the stab was either in the center of the magnet placed lengthwise under the plate or at one pole. Control plates were set away from the magnets. The plates were incubated at 30° C for 16 hours. Images of the plates were taken by using a ChemiDoc illuminator and the diameter of the bacterial circle was recorded.

It is known that flagellar motion of *Vibrio* depends on the concentration of NaCl in the medium, so another series of motility experiments was performed by varying the salt concentration in the medium. Three sets of motility plates were prepared; one with no NaCl, one with the standard NaCl and a third with twice the salt concentration. After stabbing with *Vibrio* sampled from a streaked plate, the plates were incubated at 30° C for 24 hours. Images of the plates were taken by ChemiDoc and the diameters of the circles produced by the growing, motile bacteria were also recorded. Another set of plates was exposed to magnetic fields by under the same conditions for motility experiments.

Growth experiments were conducted with the same setup illustrated in Figure 1 except that tubes with 1.0 mL of LB medium were monitored by turbidity measurements in the presence and absence of magnetic fields. The medium was prepared by innoculating frozen bacteria (*Vibrio, Pseudomonas* or *Enterobacteria*) in fresh LB medium. After growing in a shaking incubator at 30° C overnight, 5µL samples were added to 1.0 mL of LB medium. To test whether magnetic fields affected bacterial growth, control groups were incubated overnight at 30° C in the absence of magnets while experimental groups were exposed to magnetic fields applied by the neodymium magnets with north-south poles placed on either side of the tube so that the 1.0 mL culture medium was exposed directly to the magnetic field. After 16 hours of growth, 100 µL samples of each culture were mixed with 900 µL of fresh LB medium in a plastic cuvette and turbidity was measured at 500 nm wavelength with a spectrophotometer. The first experiments were performed with stationary cultures. In a second series the cultures were continuously stirred to see if vortexing affected growth rates.

## 3. Results

### Microscopy of motile species

Gram staining provided an initial indication that a magnetic field can affect the composition of mixed motile bacteria. Figure 3 shows a gram stain of the original culture and it is clear that both gram-positive and gram-negative bacteria are present, as expected.

**Figure 3.**
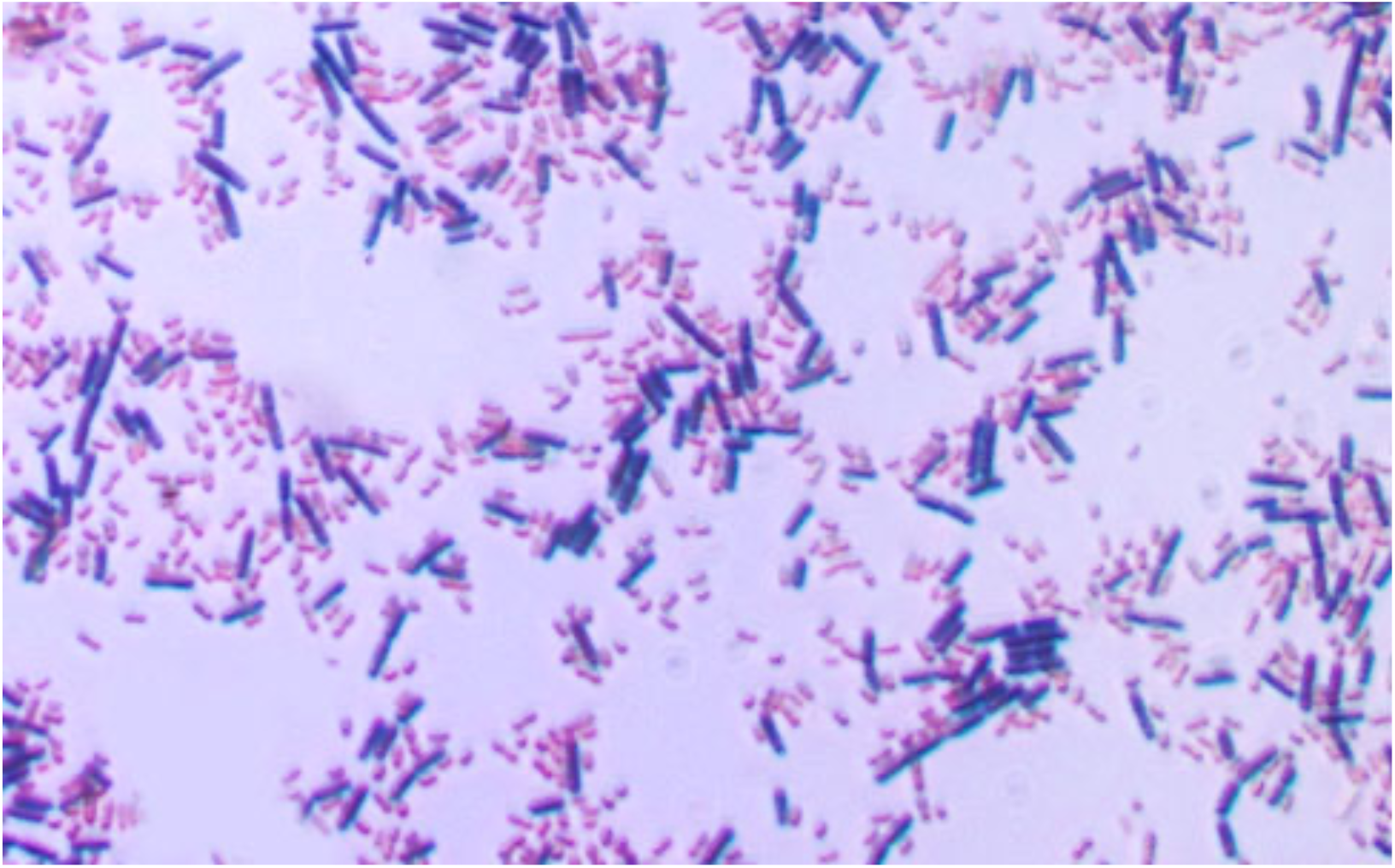
The starting culture is a mixed population of both gram-positive (blue) and gram-negative (pink) bacteria.

However, the composition of motile bacteria that had migrated through the magnetic field was significantly different from those that migrated in the absence of a field (Figure 4). Gram positive and negative species were present in the controls, but the populations of motile species that swam through the magnetic field were dominated by gram-negative species. Figure 4 shows another characteristic of the motile populations that passed through a magnetic field, which is that the number of bacteria appeared to be sparser then the controls. In order to make a quantitative determination whether this was correct, control and magnet images were subjected to a significance test for sample means. Random fields of control and magnetic samples were taken and the number of bacteria in each field was counted using ImageJ software. The control fields had a mean of 51 cells with a standard deviation of 18 while the magnetic samples had 26 cells with a standard deviation of 11. The difference was significant with a p-value less than 0.01, indicating that only half as many motile bacteria were able to navigate through the magnetic field.

**Figure 4.**
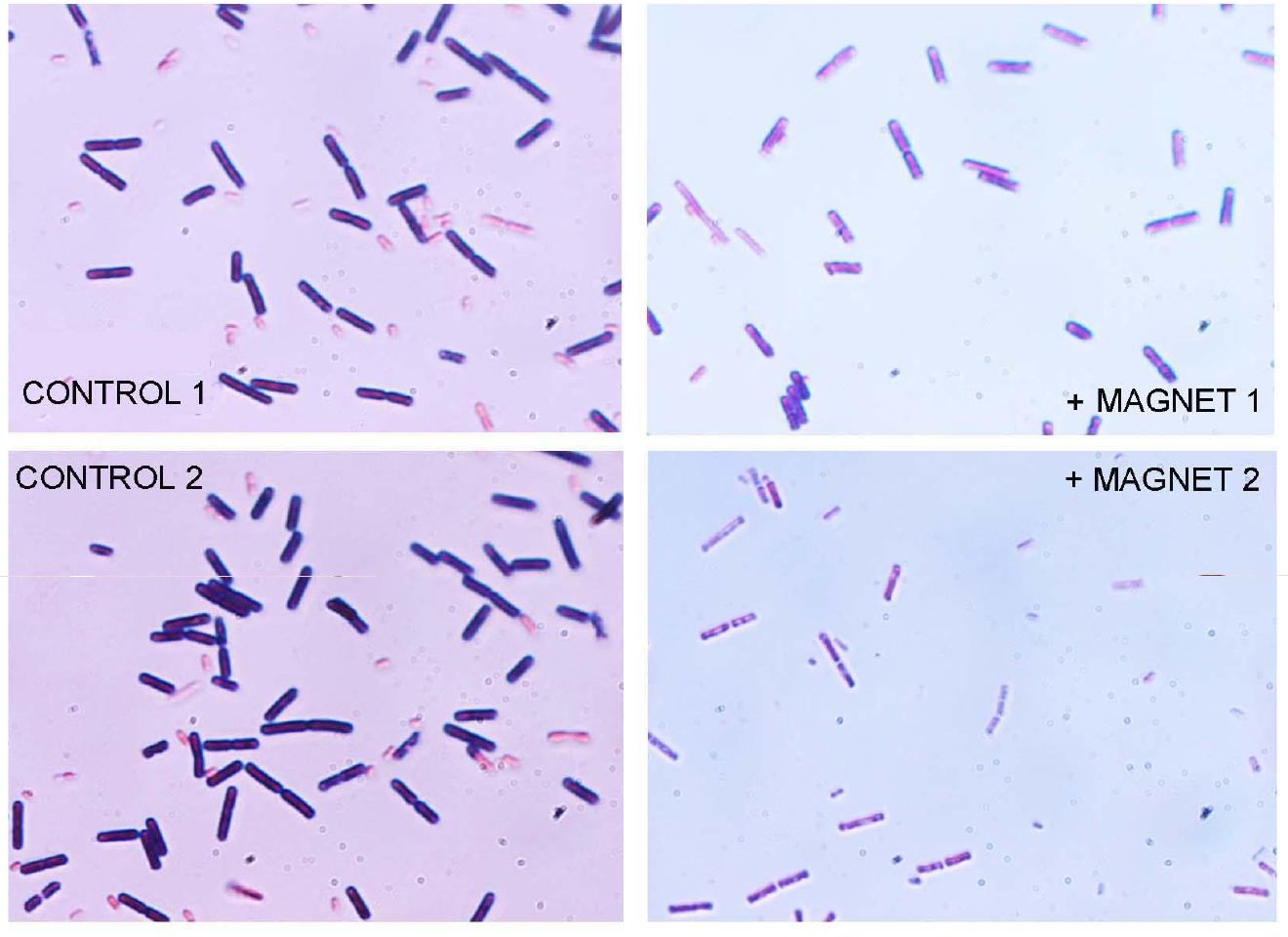
Microscopic analysis showed the filtration effects of the magnetic field. The experiment was run multiple times and two typical results are illustrated. Samples exposed to the magnetic field were sparser and contained mostly gram-negative bacteria. Original magnification 1000X.

### 16s rRNA analysis: Species diversity

The three most abundant bacterial families in the samples from the tops of the tubes were ordinary soil bacteria classified as alphaproteobacteria, gammaproteobacteris and firmicutes. Examples included *Enterobacteriaceae* species, *Escherichia coli, Comamonas testosteroni* and *Pseudomonas*. These are all motile species, as expected. Furthermore, the species that swam through a magnetic field were gram-negative which confirmed the results of microscopic analysis.

Some bacterial families that were dominant in the control groups were apparently filtered out by the magnetic field. These include *Clostridia, Erysipelotrichia*, and some species of *Lactobacillus*. All of these families are gram-positive bacteria, again confirming the microscopic observation that a magnetic field has a filtering effect on gram-negative and gram-positive species.

The Venn diagrams in Figure 5 summarize the results of the study in terms of overlapping taxonomic units, or OTUs. These do not refer to species, but instead more generally to sequences in the 16S rRNA that are revealed to either overlap or not [5]. If there were no effects, the circles would overlap completely. As expected for soil samples taken at different times of the year (dry soil in August 2020 and moist soil in April 2021) the species composition varied, but in both cases the presence of a magnetic field altered the composition. For instance, in the 2020 cultures 89 sequences were exclusively present in the control samples, 51 were exclusively present in the magnetic samples, and 56 overlapped. In the 2021 experiment 27 were exclusively in the control samples and 83 exclusively in the magnetic samples with 42 overlapping. This result further strengthens the conclusion that a magnetic field effectively filters motile bacterial species into two populations having significant compositional differences.

**Figure 5.**
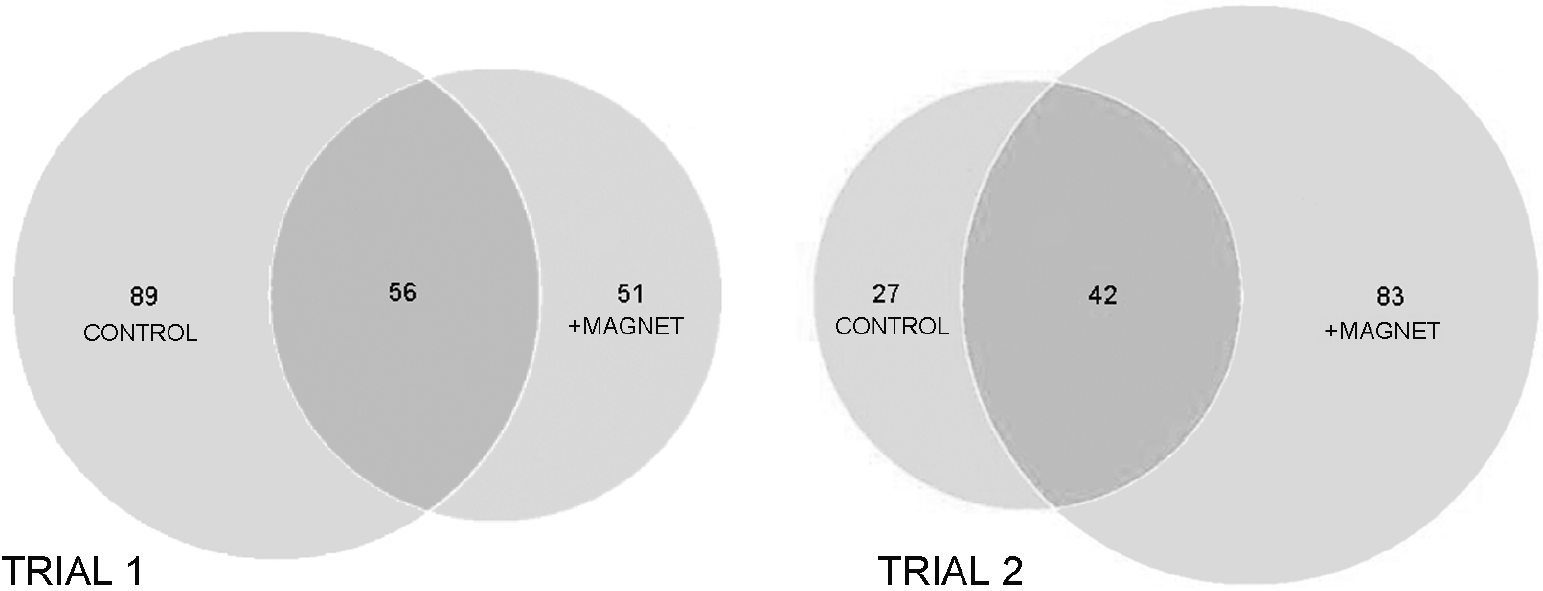
Venn diagram showing distinct differences in the populations of control motile species and those that swam through magnetic fields. Trial 1 was performed with soil samples taken in August 2020 and Trial 2 with soil samples taken in April 2021.

To summarize the results, both microscopy and 16S rRNA analysis demonstrated that a neodymium magnetic field filters out the majority of gram-positive bacteria, leaving mostly gram-negative motile species to dominate the samples. Statistical analysis performed on a series of microscopy images also indicated that the number of bacteria present in magnetic samples was less than control groups by a statistically significant amount at the p < 0.01. In two repeats that analyzed species composition by 16S rRNA, the results confirmed the filtering effect of a magnetic field.

### Potential effects of magnetic fields on known species of bacteria

It was important to determine whether magnetic fields have a general effect on bacterial motility by influencing the ionic current flowing through the flagellar motor. We chose to test three bacterial species in this regard: *Pseudomonas* and *Enterobacter* use a proton current to drive the motors, while *Vibrio* uses a sodium current. The first two were cloned from highly motile bacteria that were present in the soil bacteria samples and identified by 16S RNA analysis, while *Vibrio* samples were taken from cultures maintained in the laboratory.

Figure 6 shows a panel of growth and motility circles in petri plates 16 hours after stab inoculations of the cultures in the center of the dishes. If the magnetic fields affected bacterial motility we would expect the circles to be distorted, but there was no discernable effects on the shape of circles, although we occasionally observed what appeared to be an altered motility emerging from the *Vibrio* circle (Figure 6, upper right) but not in *Pseudomonas* or *Enterobacter*. To test whether this was related to the presence of a magnetic field, the experiment was repeated four times with *Vibrio* but the effect was not consistently observed.

**Figure 6.**
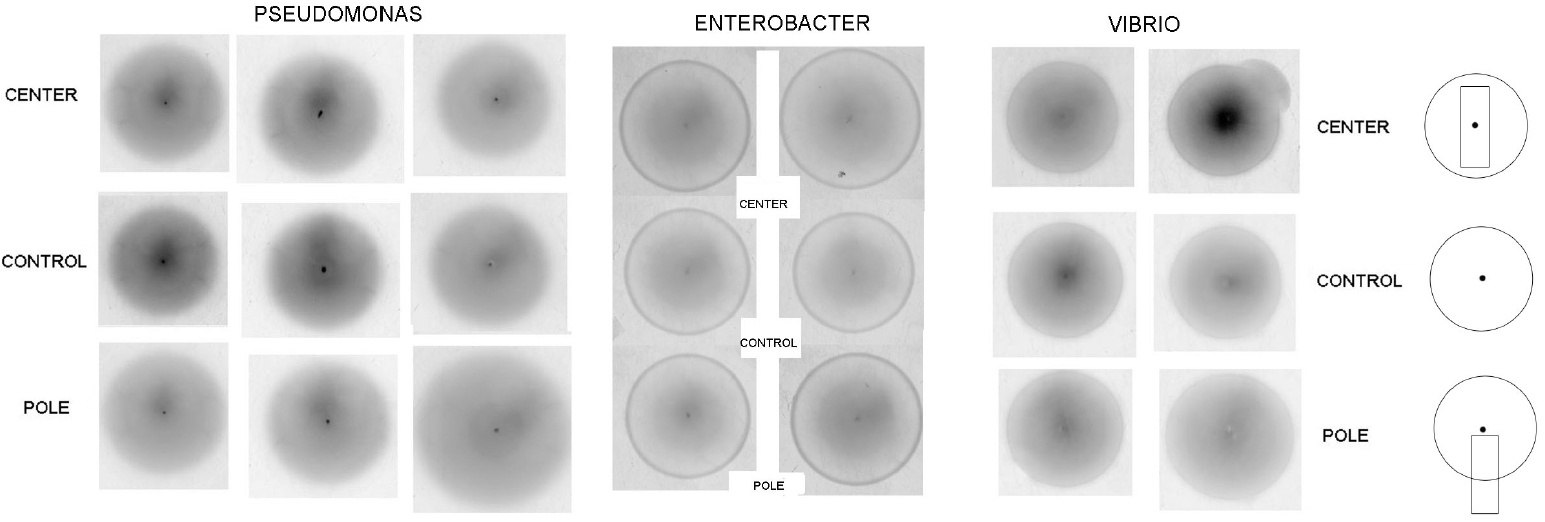
Motility assays of three bacterial species in the presence or absence of magnetic fields. Three experimental repeats are shown for *Pseudomonas*, and two for *Enterobacter* and *Vibrio*. Diagram on the right shows how petri plates were placed over the magnets.

Figure 7 compares tests of magnetic fields on growth of *Enterobacter, Pseudomonas* and *Vibrio* species in LB media. The units are optical density (OD) measured by a spectrophotometer after 16 hours of growth in 1.0 mL of LB culture medium as described in the Methods section. The results indicate that magnetic fields do not affect the growth of three known species of bacteria.

**Figure 7.**
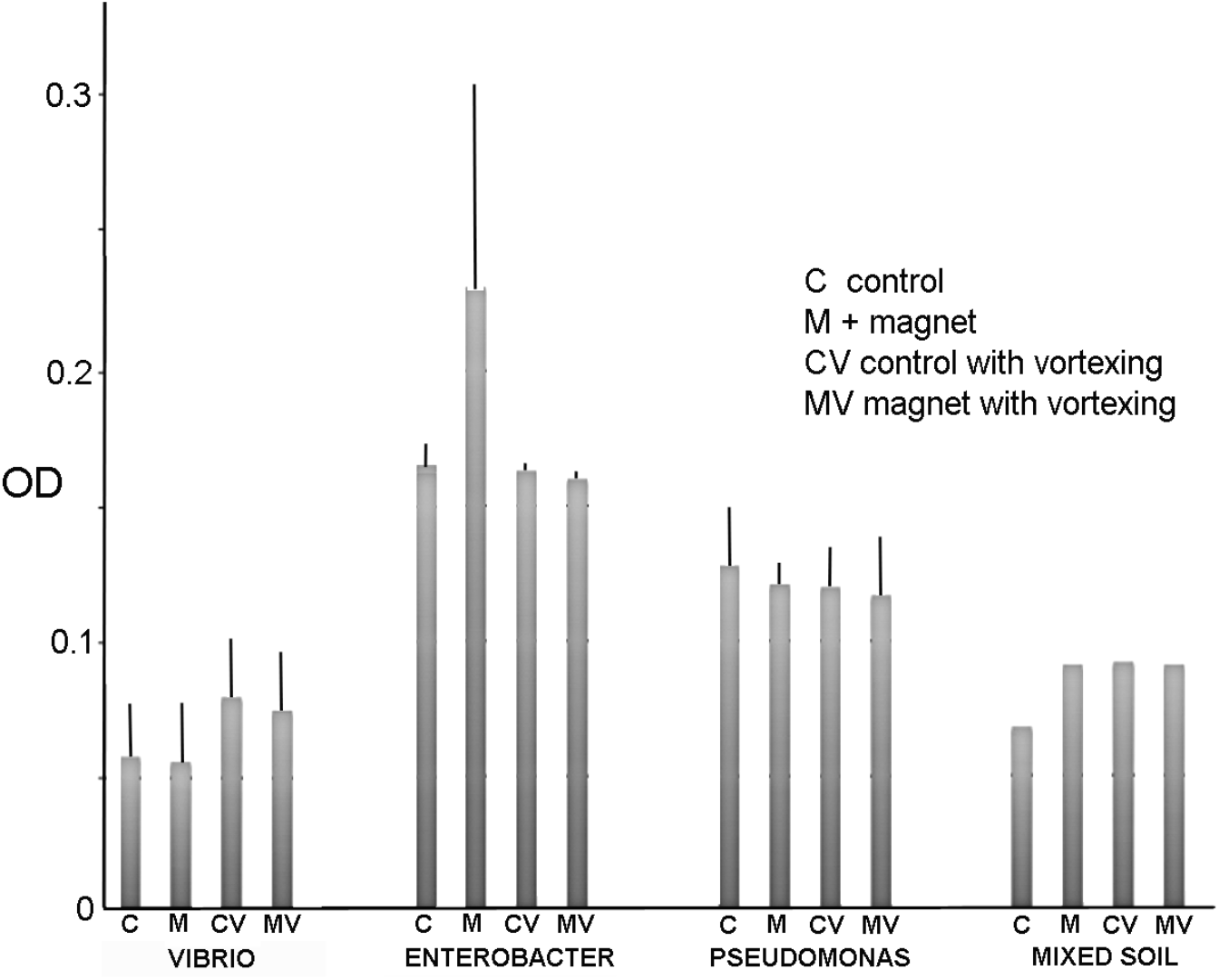
Results for growth experiments monitored by turbidity in the presence or absence of a magnetic field (mean +/S.D). The magnetic field was applied to 1.0 mL culture medium with the north and south poles of two magnets placed on either side of the medium. The results were collected after 16 hours and each set was either stationary or vortexed as indicated. OD was measured to compare the growth rates. No statistically significant differences were observed. *Vibrio* n = 6; *Enterobacter* n = 3; *Pseudomonas* n = 4; mixed soil bacteria n = 1.

## 4. Discussion

Earlier studies observed that static magnetic fields have a variety of effects on pure cultures of bacterial species [6-9]. To our knowledge, a filtering effect on mixed cultures of soil bacteria has not been previously reported. Magnetic filtering may be a selective factor in the evolution of magnetotaxis as will be discussed later.

### Force of a magnetic field acting on ionic currents

The proton current through the flagellar motor is driven by a protonmotive force equivalent to ∼170 mV generated by the electron transport system of the bacterial membrane [10]. The voltage is developed by a constant outward pumping of protons, while other protons flow back into the cell through the flagellar motors and the ATP synthases that generate ATP. Although ∼100,000 protons per second pass through a flagellar motor, the path of an individual proton is not like the path of an electron through a copper wire. Instead, there is a transduction of electrical energy to mechanical energy so that the current is doing work against a resistance. The mechanism is not understood in detail, but does involve the protonation and deprotonation of an aspartic acid residue near the cytoplasmic side of the motor [11]. The transport process leading up to the aspartate most likely involves protons jumping along hydrogen bonded chains of water rather than a simple current of protons. However, the net result is like a current in that it involves vectorial motion of charged species through space. In contrast, sodium currents involve a voltage-driven flux of the hydrated ions through the motor.

This complicated process can be simplified if we assume that a magnetic field influences the motion of ions as though they were an ordinary electrical current. The effect can be calculated from the equation below in which the force acting on an ion equals the charge multiplied by the velocity of the ion and magnitude of the magnetic field [12]:

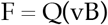

where F is the force in newtons, Q is the electrical charge on the electron or ion, v is the velocity of the ion or electron in the current and B is the amplitude of the magnetic field in teslas (T).

For instance, the electrical charge of a proton is 1.602 × 10^−19^ coulombs and the strength of the magnetic field of our neodymium magnets is 1.25 T. The velocity of a proton in the current can be calculated from the proton current which has been measured to be 78,000 protons per second flowing through a pore in a 5 nm thick membrane [4]. It follows that the velocity of one proton in the current is 5 X 10^−4^ meters per second. Because the bacteria have no particular orientation in space, protonic current will be affected only when the vector is perpendicular to the magnetic field in three-dimensional space. The cross product can be accounted for by multiplying by sin(90). Plugging in the values, the force acting on a single proton is 1.6 X 10^−19^ X (5 X10^−4^) (1.25) (0.89) = 8.9 X 10^−23^ newtons. To calculate the total force acting on a flagellar motor of a bacterium, this value must be multiplied by 10^5^ protons per second, which amounts to 8.9 X 10^−18^ newtons. For comparison, the torque generated by the *E. coli* rotor is ∼6 X 10^−18^ newton meters [13].

At first glance, it might seem that the force generated on the proton current by the magnetic field could be significant, particularly because the force acts on a very small flagellar mass. Furthermore, mixed populations of soil bacteria have varying sizes and numbers of flagella, and this could account for the filtering effect of a strong magnetic field. We reported here that gram negative species are less affected by the magnetic field and therefore dominate the population that reaches the top of the fresh medium in the tubes. However, the fact that we did not observe any influence of strong magnetic fields on motility or growth of three known bacterial species indicates that a magnetic field does not directly cause filtering of mixed soil bacteria species by imposing a physical force on the ionic current which in turn slows flagellar rotation. The filtering effect we observed must therefore be due to another causal mechanism.

One possibility is that some species of bacteria have an electrical dipole and are therefore given a vector when they move in a magnetic field. For instance, cyanobacteria are evolutionary ancestors of chloroplasts in plants, and it is known that chloroplasts become oriented so that their thylakoid stacked membranes become oriented by electrical or magnetic fields [14]. If so, this could account for the filtering effect reported here. For instance, gram positive bacteria may become aligned with the magnetic field in such a way that the vertical vector of their motion is reduced in magnitude. If gram negative bacteria lack this alignment, they would be free to swim upward through the field and accumulate near the top of the growth medium.

### Implications of magnetic filtering on bacterial evolution

Magnetic fields are not usually considered to be selective factors in evolution. The reason is that the Earth’s natural magnetic field is very weak, with surface values ranging from 25 to 65 mT, but magnetite minerals called lodestones have an intrinsic magnetic field strength thousands of times greater, in the range of 10 - 30 mT. Magnetite (Fe_3_O_4_) and hematite are the main components of iron ore on today’s Earth. Lodestones are quite rare because they are most likely produced when lightning strikes iron ore deposits [15].

A few species of motile bacteria are magnetotactic such that the direction of their motion is aligned with the magnetic field of the Earth [16]. They do this by precipitating nodules of magnetite in their cytoplasm which act as a compass needle to direct their motion. Precipitation of magnetite nodules and magnetotaxis are complex processes and were unlikely to have emerged spontaneously in a single generation but instead evolved through stages as bacteria became increasingly motile over evolutionary time. Magnetotactic bacteria are gram negative, and an interesting possibility is that their ancestral species may have originated in association with magnetic banded iron minerals that tended to exclude gram positive species.

## Author Contributions

Conceptualization, D.D. and M.N.; methodology, M.N. and E.K.; investigation, M.N. and E.K.; resources, D.D.; writing—original draft preparation, D.D.; writing—review and editing, D.D., M.N. and E.K. supervision, D.D.; project administration, D.D. All authors have read and agreed to the published version of the manuscript.

## Acknowledgments

The authors thank Prof. Fitnat Yildiz for providing bench space and supplies for this research, and Michael Trebino for his helpful advice related to culturing bacterial species.

